# Altered Developmental Trajectories in Intrinsic Function between Default, Salience, and Executive Networks in High-Functioning Autism

**DOI:** 10.1101/252320

**Authors:** Liu Yang, Xiao Chen, Xue Li, Yang-Qian Shen, Hui Wang, Jing-Ran Liu, Ting Mei, Zhao-Zheng Ji, Yan-Qing Guo, Fei-Fei Wang, Ling-Zi Xu, Xin-Zhou Tang, Zeng-Hui Ma, Li-Qi Wang, Jing Liu, Qing-Jiu Cao, Chao-Gan Yan

**Affiliations:** Peking University Sixth Hospital/Institute of Mental Health, National Clinical Research Center for Mental Disorders (Peking University Sixth Hospital), Beijing, China; Key Laboratory of Mental Health, Ministry of Health, Peking University, Beijing, China; CAS Key Laboratory of Behavioral Science, Institute of Psychology, Beijing, China; Department of Psychology, University of Chinese Academy of Sciences, Beijing, China; Magnetic Resonance Imaging Research Center, Institute of Psychology, Chinese Academy of Sciences, Beijing, China; Department of Child and Adolescent Psychiatry, NYU Langone Medical Center School of Medicine, New York, NY, USA

**Keywords:** autism, central executive network, default mode network, developmental trajectory, salience network

## Abstract

Although many studies have focused on abnormal patterns of brain functional connectivity in Autism spectrum disorder (ASD), one important factor, the developmental effect of brain networks was largely overlooked. To clarify the abnormal developmental trajectory of brain functional connectivity in ASD, we focused on the age-related changes in three “core” neurocognitive networks: default mode network (DMN), salience network (SN) and central executive network (CEN, also divided into left and right CEN, i.e., lCEN and rCEN). The development of intrinsic functional connectivity (iFC) within and between these networks were analyzed in 107 Chinese participants, including children, adolescents, and adults (54 patients with ASD and 53 typically developed (TD) participants; ages 6-30 years). We found that diagnosis-related distinctions in age-related changes suggest three maturation patterns in networks’ or nodes’ iFC: delayed (iFC between SN and rCEN), ectopic (iFC between SN and DMN, and iFC between posterior cingulate cortex (PCC) and right anterior insula/dorsal anterior cingulate cortex (dACC)), and failure maturation (iFC between dACC and ventral medial prefrontal cortex). Compared with age-matched TD participants, ASD patients in children and adolescents exhibited hypo-connectivity, while that in adults showed hyper-connectivity. In addition, an independent verification based on Autism Brain Imaging Data Exchange (ABIDE) datasets confirmed our findings of developmental trajectories in ASD group, which also showed unchanged functional connectivity with age between DMN and SN and increasing iFC between rCEN and SN. The conspicuous differences in the development of three “core” networks in ASD were demonstrated, which may lead a nuanced understanding towards the abnormal brain network maturation trajectory of autism.

## 1. Introduction

Autism spectrum disorder (ASD) is known as a heterogeneous neurodevelopmental disorder characterized by core deficits in communication and social interaction, along with restrictive interests and stereotyped and repetitive behavior ^1^, but its etiology and pathogenesis remain unclear. Since 1995, Biswal and colleagues observed synchronized spontaneous blood oxygen level-dependent (BOLD) fluctuations during rest in the motor system ^2^, resting-state functional magnetic resonance imaging (fMRI), as a mean for measuring the intrinsic functional connectivity (iFC), has been widely applied to explore the brain mechanisms in autism. Although many studies have provided evidence that ASD is a “disorder of abnormal brain connectivity” ^3, 4^, their findings are not consistent. Some studies suggested a decreased iFC between or within brain networks in subjects with autism ^5–11^, some showed increased iFC ^7, 12–14^, while others indicated a combination of both ^15–17^.

These mixed results may be contingent on the heterogeneous nature of ASD studies, such as methodological choices (low-pass filtering, global signal removal, task regression, seed selection, and field of view) ^18^, inclusion criteria, sample sizes, analysis techniques, etc. However, another key factor could be that many studies overlooked the developmental effect of brain networks in ASD. Since the developmental trajectory of ASD is variable with early life attack ^19^, subjects’ age ranges and developmental stages should be taken into account ^20^. In recent years, studies on typical brain maturation have demonstrated that iFC changed over age significantly ^21–26^ and a developmental pattern of “local to distributed” has been proposed ^23, 27^. Meanwhile, lots of ASD related studies also began to explore the brain developmental trajectories. For example, age-related changes in brain size ^28^, cortical thickness (Wallace et al. 2010), local or whole grey matter volumes ^29, 30^, and white matter microstructure ^31, 32^, have been observed to be different in ASD subjects than in controls. For functional connectivity, studies on task modulated brain function showed atypical age-related changes in the networks functioning of self-related empathizing ^33^ and face-processing ^34^. Also, resting-state fMRI has been applied to investigate the earlier stages of ASD development in the scope of iFC and functional brain networks ^35, 36^. Specifically, in domain-specific regions, age-related changes have been explored in the striatum ^37^, and superior temporal sulcus ^38^. In brain networks, default mode network (DMN) has been found to have a different developmental trajectory in ASD compared to controls ^39^. Within-and between-networks, atypicalities of FC in ASD were not uniform across the lifespan, demonstrated by the studies that used an entirely data-driven approach in age-stratified cohorts of ASD subjects and healthy controls ^38, 40^. Thus, converging all the results above, it is clear that exploring the brain mechanism of autism from a developmental perspective combined with age effects can lead to a more nuanced understanding about the atypicalities of FC in ASD.

In 2011, Menon ^41^ proposed a triple network model of major psychopathology, which suggested that aberrant intrinsic organization of within and between network connectivity were characteristics of many psychiatric disorders. The model consists of the default mode network (DMN), the salience network (SN), and the central executive networks (CENs). The DMN is implicated in self-referential and stimulus-independent memory recall ^42^. The CENs are robustly associated with inhibition, attentional control, working memory, set shifting, monitoring behaviors, and other higher cognitive processes ^43^. The SN has been shown to play a role in detecting, integrating and filtering the external and internal information (e.g. attentional and self-related mental processes) ^43^, and mediating the interactions between DMN and CEN ^44^. The triple network model has been widely applied to illuminate the brain mechanisms of multiple disorders, including Alzheimer’s disease ^45^, schizophrenia ^46, 47^, depression ^48^, etc. Recently, Abbott and colleagues investigated the iFC of these networks in 37 ASD and 38 typically developed (TD) participants ^49^, which was the first research focusing on the three “core” neurocognitive networks in autism. In this study, they found network-specific patterns in whole-brain analyses and associations between reduced SN integrity and sensory and socio-communicative symptoms. The authors also found DMN-lCEN connectivity demonstrated age-by-group interaction, however, due to the limited age range (8∼17) and samples size (37 vs. 38), the authors couldn’t examine the altered maturation patterns from child to adult of these networks in ASD.

Given the potential value of the brain functions’ development and the triple network model in explaining aberrant cognitive processes and symptoms of mental disorders, we designed this study to examine the iFC within and between DMN, SN and bilateral CENs in ASD and TD participants, from a developmental perspective (childhood to adulthood). According to previous analysis ^40, 49, 50^, we predicted that aberrant within-and between-network connectivity would be observed in ASD relative to TD in age-stratified cohorts. However, we could not specify directional hypotheses for the developmental trajectory because of limited existing evidence on it, particularly the connectivity within or between networks in ASD and their diagnostic group-by-age interactions. Therefore, an exploratory study was conducted to provide a first detailed delineation of the triple network’s age-related changes in ASD and differences relative to TD. To our knowledge, this study is the first to access the maturation patterns of iFC of the triple-network model in ASD. In order to explore the age-related effects thoroughly, we enrolled relatively “pure” subjects to ensure the samples’ homogeneity that all ASD subjects should meet the criteria of autistic disorder in DSM-IV, full IQ≥70, right-handedness, no other comorbidities, and no current psychoactive medications. Furthermore, the same analysis was performed by taking the samples from a public dataset, the Autism Brain Imaging Data Exchange (ABIDE; http://fcon_1000.projects.nitrc.org/indi/abide) ^7^ with equal screening conditions to verify this study’s reproducibility.

## 2. Materials and Methods

### 2.1 Participants

Two datasets were included in this study: PKU dataset was collected by ourselves, and the ABIDE dataset was from the public. For the PKU dataset, 87 ASD patients between 6 and 27 years old were recruited from an outpatient clinic in the Institute of Mental Health, Peking University, China. Fifty-nine TD participants between 6 and 30 years old were recruited through advertising and word of mouth. All participants were right-handed and had an intelligence quotient (IQ) over 70 evaluated with either the Chinese-Wechsler Intelligence Scale for Children (C-WISC) ^51^ or the Wechsler Adult Intelligence Scale-Revised in China (WAIS-RC) ^52^ depending on their age. The ASD patients’ diagnoses were established clinically by experienced psychiatrists in our institute according to the criteria of autistic disorder in *Diagnostic and Statistical Manual-IV* (DSM-IV) (Rett’s disorder, childhood disintegrative disorder, Asperger’s disorder and pervasive developmental disorder NOS were excluded). The severity of symptoms was evaluated by a trained and qualified clinician using the Autism Diagnostic Observation Schedule (ADOS) ^53^. Same exclusion criteria were applied to both ASD and TD groups that participants with major physical or neurological diseases, current or previous psychiatric diagnoses (other than the autism spectrum disorder in ASD group), psychotropic medication prescriptions, or a history of head injury, alcohol or drug abuse were not considered in this study. The participants not suitable for magnetic resonance imaging screening, such as that having cardiac pacemakers, insulin pumps, artificial heart valves, or other metalwork in vivo, were excluded. Twelve ASD patients and 2 TD participants were excluded due to big head motion: with mean frame-wise displacement (FD) ^54^ > 0.48 (corresponding to 2 SDs above the whole-sample mean). Twenty-one ASD patients and 4 TD participants were excluded to match the gender between the two groups. Finally, 54 ASD patients and 53 TD participants who matched for gender, age, full-scale IQ level and mean FD were included in the final statistical analysis (**Table 1**). Considering this study’s nature of cross-sectional analysis and its potential sampling bias related to age differences, we investigated the relationships between age and the severity of ASD symptoms, which were evaluated by the ADOS scales involving various aspects (communication, social and restricted repetitive behavior). No significant age effect was shown in any ADOS examination within the ASD group (*p*>0.05).

**Table 1.** Demographic information

| | ASD (n=54) | TD (n=53) | $t/\chi^2/z$ | $p$ |
| --- | --- | --- | --- | --- |
| Gender (male:female) | 45:9 | 40:13 | 1.01 | 0.314 |
|  | Mean±SD | Mean±SD |  |  |
| Age | 13.56±5.04 | 14.51±6.06 | -0.89 | 0.378 |
| Full-scale IQ | 110.70±15.50 | 115.00±11.40 | -1.64 | 0.103 |
| Verbal IQ | 114.60±17.70 | 116.60±12.90 | -0.67 | 0.505 |
| Performance IQ | 102.80±15.10 | 109.90±12.20 | -2.68 | 0.009 |
| ADOS<sup>a</sup> |  |  |  |  |
| Total (Communication+ Social) | 10.70±3.78 | - | - | - |
| Communication | 2.95±1.65 | - | - | - |
| Social | 7.76±2.36 | - | - | - |
| RRB | 1.00<sup>b</sup> | - | - | - |
| Head motion (mean FD) | 0.093<sup>b</sup> | 0.059<sup>b</sup> | 1.75 | 0.066<sup>c</sup> |

*Note:* ASD, autism spectrum disorder; TD, typical development; SD, standard deviation. IQ: intelligence quotient; ADOS, autism diagnostic observation schedule; RRB, restricted repetitive behaviors; mean FD, mean-frame wise displacement; <sup>a</sup>n=37.
<sup>b</sup>Median.
<sup>c</sup>Asymptotic significance was displayed using nonparametric test.

### 2.2 Ethical Considerations

The research procedure was approved by the Ethics Committee of Institute of Mental Health, Peking University with the written informed consent from all participants or their parents.

### 2.3 Data Acquisition

A GE-Discovery 3T MR750 scanner with an 8-channel head coil at the Peking University Third Hospital was utilized to collect the imaging data. Subjects completed the resting-state fMRI scanning for 8 mins. A total of 240 whole-brain functional volumes were obtained for each one by using a multi Echo-Planar Imaging (EPI) sequence. Detailed parameters were shown as follows: repetition time (TR)=2.0s; echo time (TE)=20 ms; flip angle=90; 41 slices; matrix=64×64; and voxel size=3.75×3.75×3.30mm. All participants were guided to simply rest with eyes closed but being awake during the scanning. A high-resolution T1-weighted structural image was also obtained through a magnetization-prepared gradient-echo sequence with the parameters of TR=4.78 ms, TE=2.024 ms, inversion time (TI)=400 ms, flip angle=15, 166 slices, the field of view (FOV)=240×240 mm, and voxel size=0.94×0.94×1mm.

### 2.4 fMRI Preprocessing

Unless otherwise stated, the Data Processing Assistant for Resting-State fMRI ^55^, was applied to preprocess all of the functional imaging data. This software was developed based on the platform of Statistical Parametric Mapping (SPM, http://www.fil.ion.ucl.ac.uk/spm) and the toolbox for Data Processing & Analysis of Brain Imaging ^56^. The preprocessing procedure for each subject followed the steps below: 1) the first 10 time points of the fMRI series were discarded to allow for the signal equilibration and adaptation of participants to the scanning environment. 2) The remaining 230 time-points were corrected for different signal acquisition time by taking the mid-point slice as the reference to shift other signals. 3) A six-parameter (rigid body) linear transformation with a two-pass procedure (in turn, registered to the first image and the mean image after the first realignment) was used to realign the time series of images. 4) Corrected the motion by using a 6 degrees-of-freedom linear transformation. 5) Co-registered the T1-weighted image to the mean functional image.

6) A unified segmentation algorithm ^57^ was applied to segment the transformed T1 image into three components: gray matter (GM), white matter (WM) and cerebrospinal fluid (CSF) that were referred by an SPM’s prior tissue maps. 7) Transformed individual native space to Montreal Neurological Institute (MNI) space based on the Diffeomorphic Anatomical Registration algorithm computed by the Exponentiated Lie algebra (DARTEL) ^58^. 8) Considering the head motion effects, the Friston 24-parameter model ^59^ (i.e., 6 head motion parameters, 6 head motion parameters one time point before, and the 12 corresponding squared items) was chosen to regress out this covariate based on previous findings that higher-order models can better remove head motion effects ^60, 61^. To address the residual effects of motion in group analyses, mean FD was taken as a covariate. 9) White matter signal and cerebrospinal fluid signal were regressed out from each voxel’s time course, and linear trends were included as a regressor to account for drifts in the blood oxygen level dependent (BOLD) signal. 10) Finally, all images were filtered by temporal band-pass filtering (0.01-0.1Hz) to reduce the effect of low frequency-drift and high-frequency physiological noise. Further data analysis and visualization were completed using the Data Processing & Analysis for Brain Imaging DPABI, ^56, http://rfmri.org/DPABI^.

### 2.5 Data Analysis

#### 2.5.1 Seed Selection

We selected 3-mm radius spherical seeds as the regions of interest (ROIs) within each network: posterior cingulate cortex (PCC), ventral medial prefrontal cortex (vmPFC), left and right angular gyri (lAng and rAng) for the DMN; right and left anterior insula (rAI and lAI), dorsal anterior cingulate cortex (dACC) for the SN; right posterior parietal cortex (rPPC), right dorsolateral prefrontal cortex (rdlPFC) for the rCEN; left posterior parietal cortex (lPPC), left dorsolateral prefrontal cortex (ldlPFC) for the lCEN (**Figure 1**, please see the coordinates of the seeds in **Table 2**). Of note, the ROIs and coordinates used in this study were derived from Chinese population based on a prior study ^48^.

**Table 2.** MNI coordinates for four networks’ seed regions

| Seed | x | y | z | Seed | x | y | z |
| --- | --- | --- | --- | --- | --- | --- | --- |
| **Default Mode Network (DMN)** |  |  |  | **Salience Network (SN)** |  |  |  |
| PCC | 0 | -52 | 18 | rAI | 34 | 23 | 5 |
| vmPFC | -1 | 56 | 10 | IAI | -34 | 23 | 4 |
| lAng | -48 | -67 | 34 | dACC | 2 | 34 | 25 |
| rAng | 54 | -61 | 30 |  |  |  |  |
| **Right Central Executive Network (rCEN)** |  |  |  | **Left Central Executive Network (lCEN)** |  |  |  |
| rPPC | 42 | -62 | 50 | lPPC | -33 | -70 | 50 |
| rdIPFC | 27 | 20 | 62 | ldIPFC | -27 | 20 | 58 |

*Note:* from (Zheng et al. 2015).

**Figure 1.**
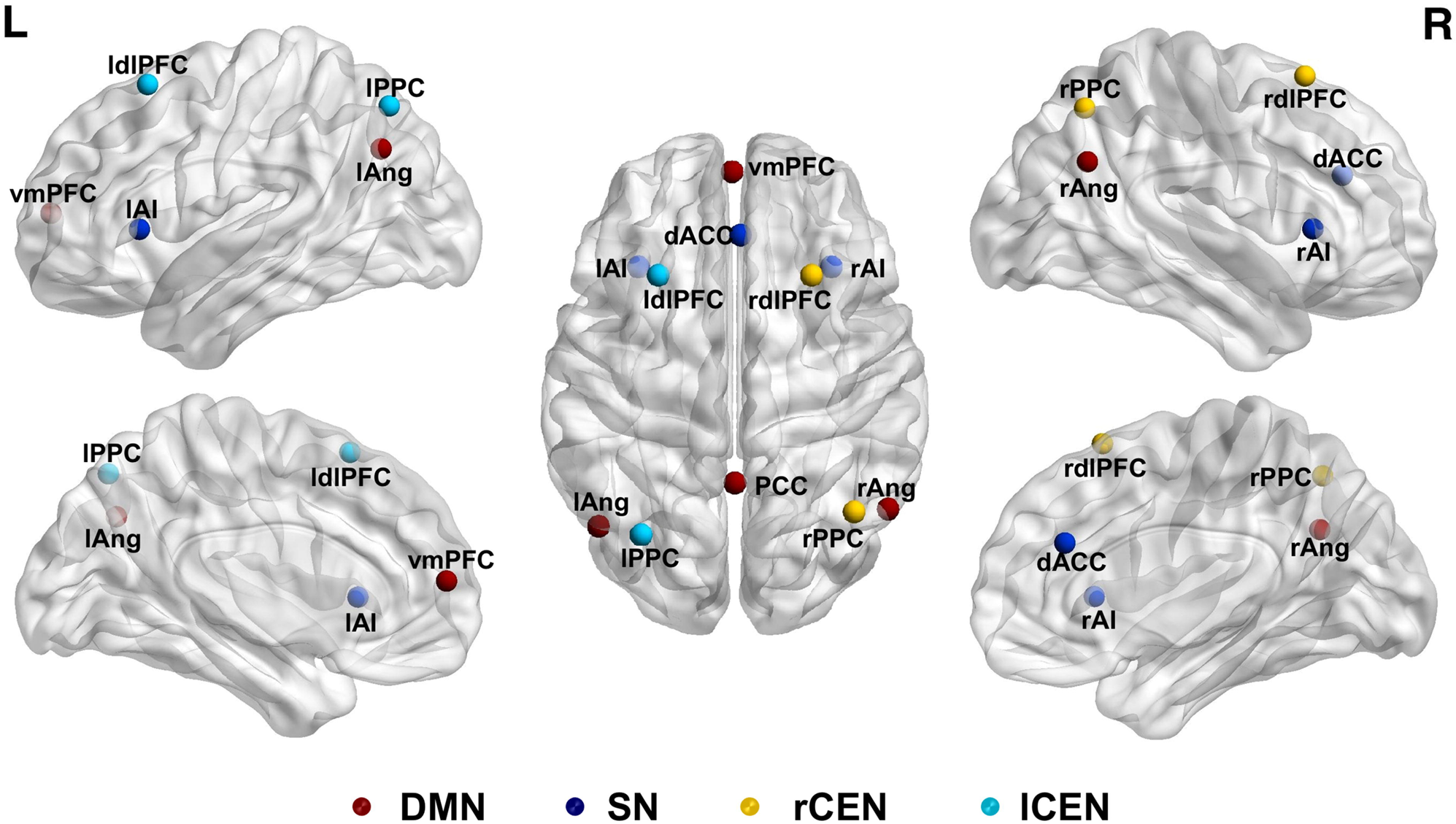
ROIs within each network: the red dots are the ROIs in the Default Mode Network (DMN); the dark blue dots are the ROIs in the Salience Network (SN); the orange dots are the ROIs in right Central Executive Network (rCEN); the light blue are the ROIs in left Central Executive Network (lCEN). Seeds consist of posterior cingulate cortex (PCC), ventral medial prefrontal cortex (vmPFC), left and right angular gyri (lAng and rAng) for the DMN; right and left anterior insula (rAI and lAI), dorsal anterior cingulate cortex (dACC) for the SN; right posterior parietal cortex (rPPC), right dorsolateral prefrontal cortex (rdlPFC) for the rCEN; left posterior parietal cortex (lPPC), left dorsolateral prefrontal cortex (ldlPFC) for the lCEN.

#### 2.5.2 Subject-level

Similar to a prior study ^49^, signal ROI time series was extracted by averaging all voxels in the given seed ROI, and then the time series in each network were calculated by averaging all ROIs time series within the network. Subsequently, the *Z*-transformed correlation coefficients (Fisher’s r to *z* transformation) were computed for all ROI and network pairs. The within-network connectivity for DMN, SN, CENs was derived separately by taking the mean of *z* values for all ROI pairings within the corresponding network. The between-network connectivities were the z values of network pairings.

#### 2.5.3 Group-level

IBM SPSS Statistics 20.0 was used for statistical analysis of the network connectivity data and the clinical scale data. Student’s *t* tests and Nonparametric test was used to determine whether there are significant group differences regarding these demographic variables.

To analyze the differences in brain network development among diagnostic groups, univariate analyses of variance procedures were conducted to identify group-by-age interaction effects. The *z* values representing within/between-network connectivity was involved as the dependent variable, while diagnosis as a fixed factor and age as a covariate were involved in this modeled diagnosis×age interaction. Gender, full-scale IQ and mean FD were controlled well in the statistical analysis. For the network showing significant group-by-age interactions, we explored the shape of developmental curves by examining a quadratic and a linear model of age that best fit the collected data in each group. Specifically, network connectivity’s z values, as the dependent variables, while age and age^2^, as predictors, were involved in linear regression analyses. Furthermore, we investigated connectivity differences across groups for three age cohorts: childhood, 6.0-11.0 years, n=36; adolescence, 11.1-15.2 years, n=36; early adulthood, 15.4-29.6 years, n=35 (**Supplementary Table 1**). To ensure an equal number of each cohort, terciles calculation was utilized to determine the age boundaries. Further analyses were conducted in ROI pairs of the networks using the same procedures described above. Finally, to assess the effect of symptom severity on connectivity, Spearman rank-order correlations were performed on the ASD group with the variables of connectivity’s *z* values and ADOS scores. Each age cohort was involved in this analysis to examine the age-specific correlations.

### 2.6 Validation Using Autism Brain Imaging Data Exchange Dataset

ABIDE is an open resting-state fMRI dataset for ASD patients and age-matched TD participants, including ABIDE I and ABIDE II ^62, 63^. To validate our results, participants in both groups were selected with the same screening criteria to guarantee the homogeneity of the diagnosis with autistic disorder and the demographic characteristics. In ABIDE II, all ASD participants were roughly diagnosed with ASD, which were not further classified into autistic disorder/Asperger’s syndrome/PDD-NOS. Thus, only ABIDE I was adopted for having a clear diagnostic category of autistic disorder according to DSM-IV-TR. Screening criteria for subjects were as follows: 1) meeting autistic disorder criteria of DSM-IV-TR with exclusion of Asperger’s syndrome and PDD-NOS; 2) 6-30 years of age; 3) full IQ≥70 that evaluated by Wechsler Scales; 4) right-handedness; 5) no other comorbidities; 6) no current psychotropic medications; 7) individuals with mean FD<2 SDs above the sample mean (0.5mm); Additionally, only sites with at least 5 participants per group after the above exclusions were included in the validation analysis. The yielded data were 96 ASDs and 101 TDs from 7 sites (**Supplementary Table 2**). The fMRI preprocessing and data analyses were same as above, plus taking sites as covariates in group analysis.

## 3. Results

### 3.1 3.1 Group Differences in Triple Networks’ Development

#### 3.1.1 Diagnostic Group-By-Age Interaction and Main Effects

For the between-network connectivities, the univariate analyses showed significant diagnostic group-by-age interactions in DMN-SN (*F*=4.332, *p*=0.040) and SN-rCEN (*F*=6.197, *p*=0.014). No main effects of group or age were found in any between-network connectivity (all *p*>0.05) (**Table 3**).

**Table 3.** Between-network connectivity and within network connectivity

| Variables | ASD | TD | Group |  | Age |  | Group×Age |  |
| --- | --- | --- | --- | --- | --- | --- | --- | --- |
|  | Mean±SD | Mean±SD | F | p | F | p | F | p |
| DMN-SN | 0.16±0.22 | 0.07±0.28 | 2.308 | 0.132 | 2.516 | 0.116 | 4.332 | 0.040* |
| DMN-rCEN | 0.26±0.24 | 0.33±0.23 | 0.676 | 0.413 | 0.652 | 0.421 | 0.025 | 0.874 |
| DMN-ICEN | 0.29±0.22 | 0.34±0.25 | 0.234 | 0.629 | 0.152 | 0.698 | 0.846 | 0.360 |
| SN-rCEN | 0.12±0.18 | 0.14±0.21 | 5.058 | 0.027 | 0.801 | 0.373 | 6.197 | 0.014* |
| SN-ICEN | 0.13±0.23 | 0.12±0.21 | 0.555 | 0.458 | 0.004 | 0.953 | 1.212 | 0.274 |
| rCEN-ICEN | 0.41±0.24 | 0.37±0.23 | 0.412 | 0.522 | 0.014 | 0.904 | 0.132 | 0.717 |
| DMN | 0.37±0.17 | 0.41±0.16 | 0.003 | 0.957 | 5.551 | 0.021* | - | - |
| SN | 0.39±0.18 | 0.38±0.16 | 1.597 | 0.209 | 1.650 | 0.202 | 1.792 | 0.184 |
| rCEN | 0.17±0.21 | 0.17±0.24 | 0.068 | 0.794 | 0.490 | 0.485 | 0.102 | 0.750 |
| ICEN | 0.26±0.22 | 0.27±0.24 | 0.830 | 0.364 | 0.537 | 0.465 | 0.797 | 0.374 |

*Note:* Univariate analysis was applied in this analysis. Dependent variables were listed in the left side in this table. The diagnosis was involved as a fixed factor, age was involved as a covariate and the diagnosis by age interaction was modeled. Gender, full IQ, and mean FD were controlled.
a. "DMN-SN, DMN-rCEN, DMN-ICEN, SN-rCEN, SN-ICEN, rCEN-ICEN" represent between-network connectivities; "DMN, SN, rCEN, ICEN" represent within-network connectivities.
b. "Group" represents main effects of groups; "Age" represents main effects of age; "Group×Age" represents diagnostic group-by-age interactions.
c. \* $p < 0.05$ .

For the within-network connectivities, no significant diagnostic group-by-age interactions or significant main effects of diagnosis was shown in the univariate analyses (all *p*>0.05). But there was a significant main effect of age in DMN (*F*=5.551, *p*= 0.021) (**Table 3**).

#### 3.1.2 Shape of Developmental Trajectories

For connectivities showing a diagnostic group-by-age interaction, namely DMN-SN and SN-rCEN, the shape of the developmental trajectories was analyzed by means of the comparison of linear and quadratic fits in diagnostic groups separately. For the DMN-SN, no fit was significant in the ASD group (linear: *t*=0.172, *p*=0.864; quadratic: *t*=0.772, *p*=0.444), while the variance in the TD group could be interpreted by a linear model (*t*=-2.494, *p*=0.016) (**Figure 2A**). Conversely, in the SN-rCEN, the variance in the ASD group (*t*=2.817, *p*=0.007) was explained by a linear model, but there was no fit found in the TD group (linear: *t*=-0.683, *p*=0.497; quadratic: *t*=-0.541, *p*=0.591) (**Figure 2C**). Additionally, a linear model could be also used to explain the variance in the DMN connectivity (*t*=3.451, *p*=0.001), which showed main effects of age. (**Figure 2E**).

**Figure 2.**
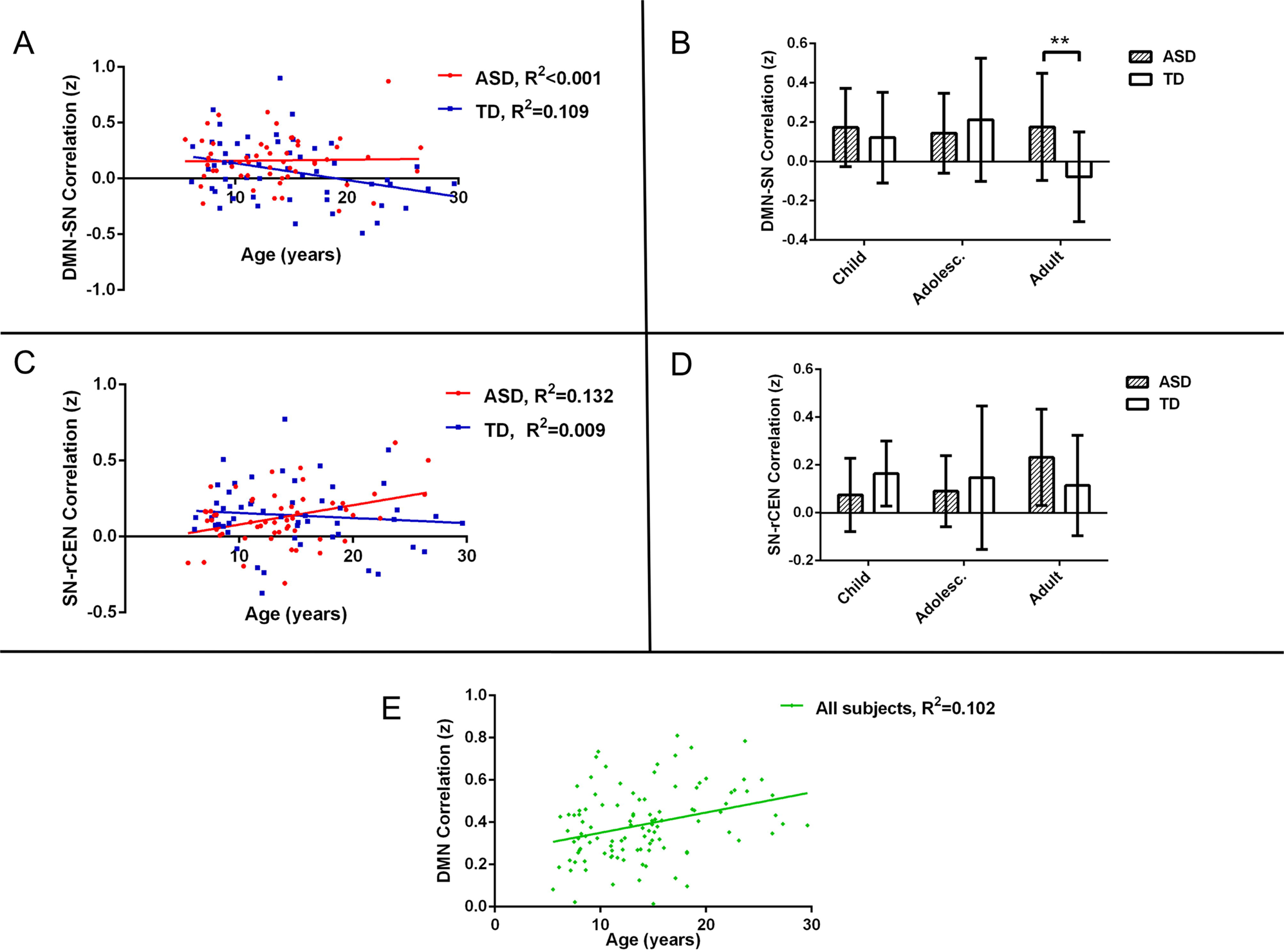
The shape of developmental trajectories and group differences in each age cohort at the level of network connectivity (z). The scatter plots and fit lines showed the diagnostic group-by-age interactions (A for DMN-SN connectivity and C for SN-rCEN connectivity) and the main effect of age (E for DMN connectivity). Diagnostic group-by-age interactions were plotted separately (red represents the ASD group and the blue represents the TD group); the main effect of age was plotted across all participants (green). The histograms showed the group differences in each age cohort corresponding to the left (B and D). Age ranges were grouped for childhood, 6.0-11.0 years; adolescence, 11.1-15.2 years; and early adulthood, 15.4-29.6 years. DMN, SN, rCEN stand for default mode network, Salience Network and right Central Executive Network, separately. \*\**p* <0.01 (uncorrected, but survived after FDR correction).

#### 3.1.3 Group Differences in Each Age Cohort

For DMN-SN and SN-rCEN revealing significant diagnostic group-by-age interactions, we divided the age into three cohorts to visualize the age effects between ASD and TD. Group comparisons within each age cohort indicated that the ASD group showed hyper-connectivity in early adulthood relative to TD group (*t*=2.999, *p*=0.005; FDR-corrected, *p*=0.015) for DMN-SN, but no significant difference in childhood and adolescence was shown (both *p*>0.05) (**Figure 2B**). For SN-rCEN, there was no significant difference found between ASD and TD in any age cohort (all *p*>0.05) (**Figure 2D**).

### 3.2 Group Differences in ROI Pairs’ Development

#### 3.2.1 Diagnostic Group-by-age Interaction and Main Effects

For the network connectivities showing diagnostic group-by-age interactions, the univariate analysis was further performed on ROI pairs in the corresponding networks. For the ROI pairs in DMN-SN, the rAI-PCC (*F*=4.083, *p*=0.046), dACC-PCC (*F*=5.788, *p*=0.018), and dACC-vmPFC (*F*=5.661, *p*=0.019) demonstrated a significant group-by-age interaction. For the ROI pairs in SN-rCEN, significant group-by-age interactions were found in the rAI-rPPC (*F*=4.125, *p*=0.045) and dACC-rPPC (*F*=7.477, *p*=0.007), and the main effect of group was found in the rAI-rdlPFC (*F*=4.799, *p*=0.031) (**Table 4**). However, only the dACC-rPPC remained significant after FDR correction (FDR-corrected, *p*=0.042).

**Table 4.** ROI pairs’ connectivity

| Variables | ASD | TD | Group |  | Age |  | Group×Age |  |
| --- | --- | --- | --- | --- | --- | --- | --- | --- |
|  | Mean±SD | Mean±SD | F | p | F | p | F | p |
| **DMN-SN** |  |  |  |  |  |  |  |  |
| rAI-PCC | 0.000±0.180 | -0.090±0.192 | 1.099 | 0.297 | 5.637 | 0.019 | 4.083 | 0.046 |
| dACC-PCC | 0.101±0.222 | 0.169±0.201 | 7.103 | 0.009 | 0.049 | 0.825 | 5.788 | 0.018 |
| dACC-vmPFC | 0.341±0.236 | 0.286±0.217 | 3.204 | 0.077 | 0.622 | 0.432 | 5.661 | 0.019 |
| **SN-rCEN** |  |  |  |  |  |  |  |  |
| rAI-rPPC | 0.064±0.178 | 0.041±0.212 | 2.688 | 0.104 | 0.259 | 0.612 | 4.125 | 0.045 |
| dACC-rPPC | 0.068±0.215 | 0.172±0.207 | 11.18 | 0.001 | 5.457 | 0.021 | 7.477 | 0.007 |
| rAI-rdIPFC | 0.067±0.177 | -0.012±0.152 | 4.799 | 0.031 | 0.024 | 0.877 | - | - |

*Note:* Univariate analysis was applied in this analysis. Dependent variables were listed in the left side in this table. The diagnosis was involved as a fixed factor, age was involved as a covariate and the diagnosis by age interaction was modeled. Gender, full IQ, and mean FD were controlled.
a. "DMN-SN, SN-rCEN" represent between-network connectivities; only ROI pairs showing significant main effects or interactions were listed above (uncorrected, only dACC-rPPC survived after FDR correction);
b. "Group" represents main effects of groups; "Age" represents main effects of age; "Group×Age" represents diagnostic group-by-age interactions.

#### 3.2.2 Shapes of Developmental Trajectories

For ROI pairs showing diagnostic group-by-age interactions, the investigation for the shapes of the developmental trajectories was implemented through the comparison of linear and quadratic fits in diagnostic groups separately. For the DMN-SN, only dACC-vmPFC showed significant linear fitting in the ASD group (*t*=2.089, *p*=0.042). In the TD group, rAI-PCC showed significant linear fitting (*t*=-2.622, *p*=0.012), while quadratic models best explained the variances in dACC-PCC (*t*=-2.384, *p*=0.021) and dACC-vmPFC (*t*=-2.147, *p*=0.037) (**Figure 3A-C**). For the SN-rCEN, only dACC-rPPC showed significant linear fitting in the ASD group (*t*=3.539, *p*=0.001), whereas no significant fit in the TD group was noticed (**Figure 3D**).

**Figure 3.**
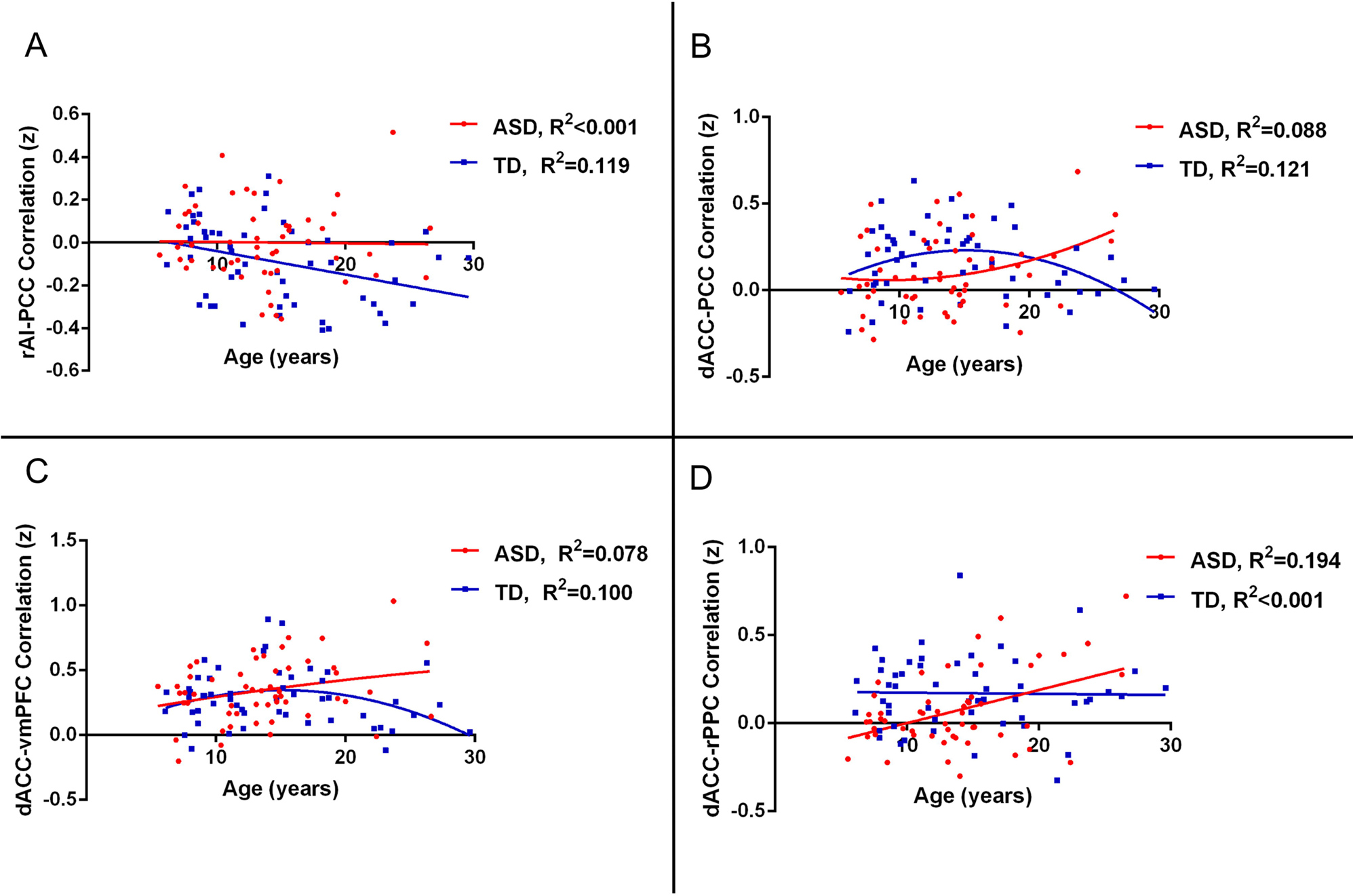
The shape of developmental trajectories at the level of ROI pairs’ connectivity (z). The scatter plots and fit lines showed the diagnostic group-by-age interactions (A for rAI-PCC connectivity; B for dACC-PCC connectivity; C for dACC-vmPFC connectivity; D for dACC-rPPC connectivity). The fit lines were plotted separately for the ASD (red) and TD (blue) group. rAI, PCC, dACC, vmPFC and rPPC stand for right Anterior Insular, Posterior Cingulate Cortex, dorsal Anterior Cingulate Cortex, ventral medial Prefrontal Cortex and right Posterior Parietal Cortex, separately.

#### 3.2.3 Group Differences in Each Age Cohort

For ROI pairs showing diagnostic group-by-age interactions, we conducted group comparisons within each age cohort. The result indicated that the ASD group showed hypo-connectivity in childhood (dACC-rPPC, *t*=-3.350, *p*=0.002; FDR-corrected, *p*=0.036) and adolescence (dACC-rPPC, *t*=-3.183, *p*=0.003, FDR-corrected, *p*=0.027; dACC-PCC, *t*=-2.292, *p*=0.028, FDR-corrected, *p*>0.05), while hyper-connectivity was found in early adulthood (rAI-PCC, *t*=4.029, *p* <0.001, FDR-corrected, *p* <0.05; dACC-vmPFC, *t*=2.823, *p*=0.008, FDR-corrected, *p*>0.05) (**Figure 4A-D, Figure 5**) relative to the TD group. For the main effects of group, the ASD group showed hyper-connectivity in rAI-rdlPFC across the whole ages (*t*=2.449, *p*=0.016, FDR-corrected, *p*>0.05).

**Figure 4.**
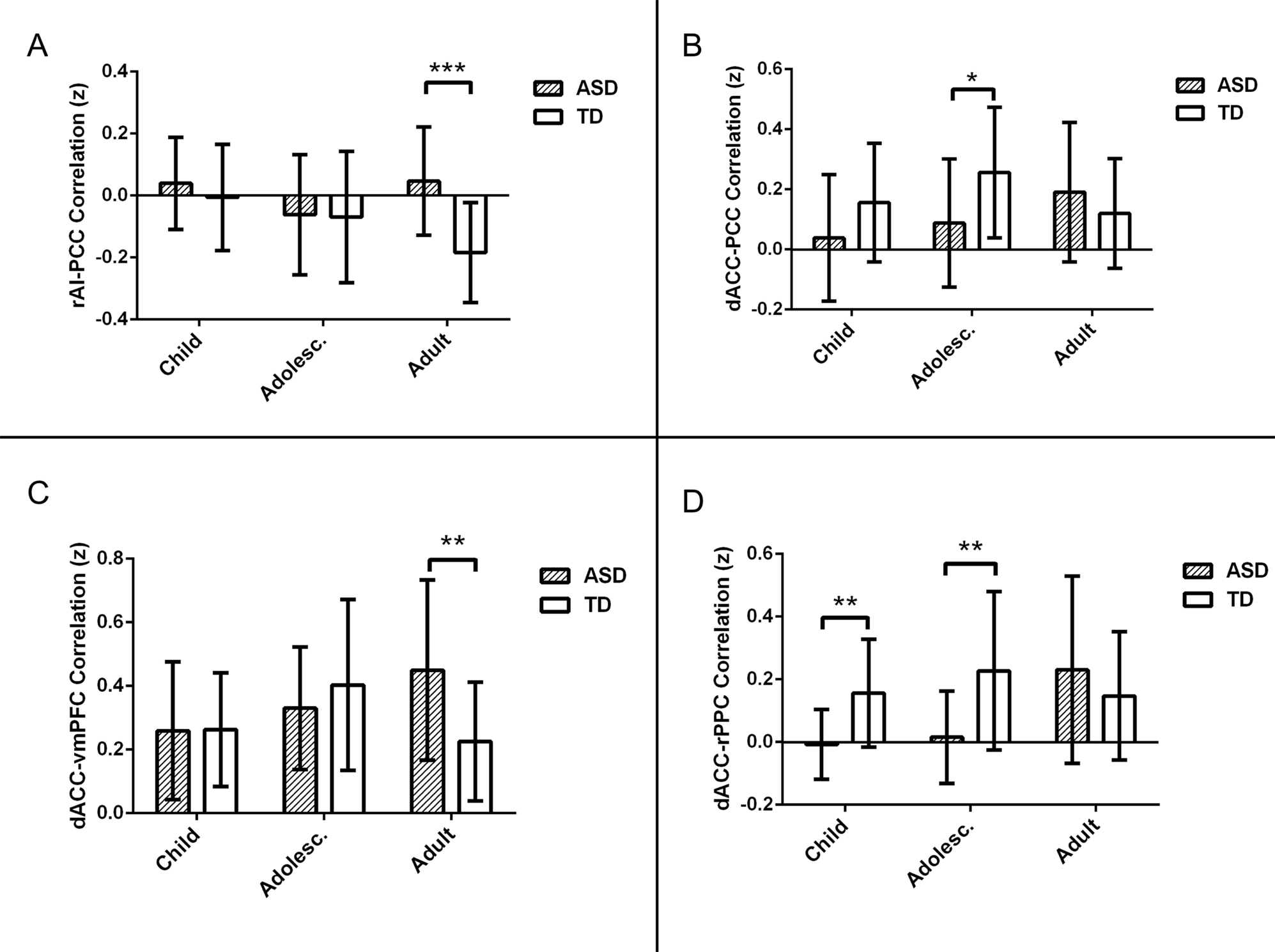
Group differences in each age cohort at the level of ROI pairs’ connectivity (z). The bar plots showed the group differences in each age cohort (A for rAI-PCC connectivity; B for dACC-PCC connectivity; C for dACC-vmPFC connectivity; D for dACC-rPPC connectivity). Age ranges were grouped for childhood, 6.0-11.0 years; adolescence, 11.1-15.2 years; and early adulthood, 15.4-29.6 years. \**p* <0.05, \*\**p* <0.01, \*\*\**p* <0.001 (uncorrected, rAI-PCC in adulthood, dACC-rPPC in childhood and adolescence survived FDR correction). rAI, PCC, dACC, vmPFC and rPPC stand for right Anterior Insular, Posterior Cingulate Cortex, dorsal Anterior Cingulate Cortex, ventral medial Prefrontal Cortex and right Posterior Parietal Cortex, separately.

**Figure 5.**
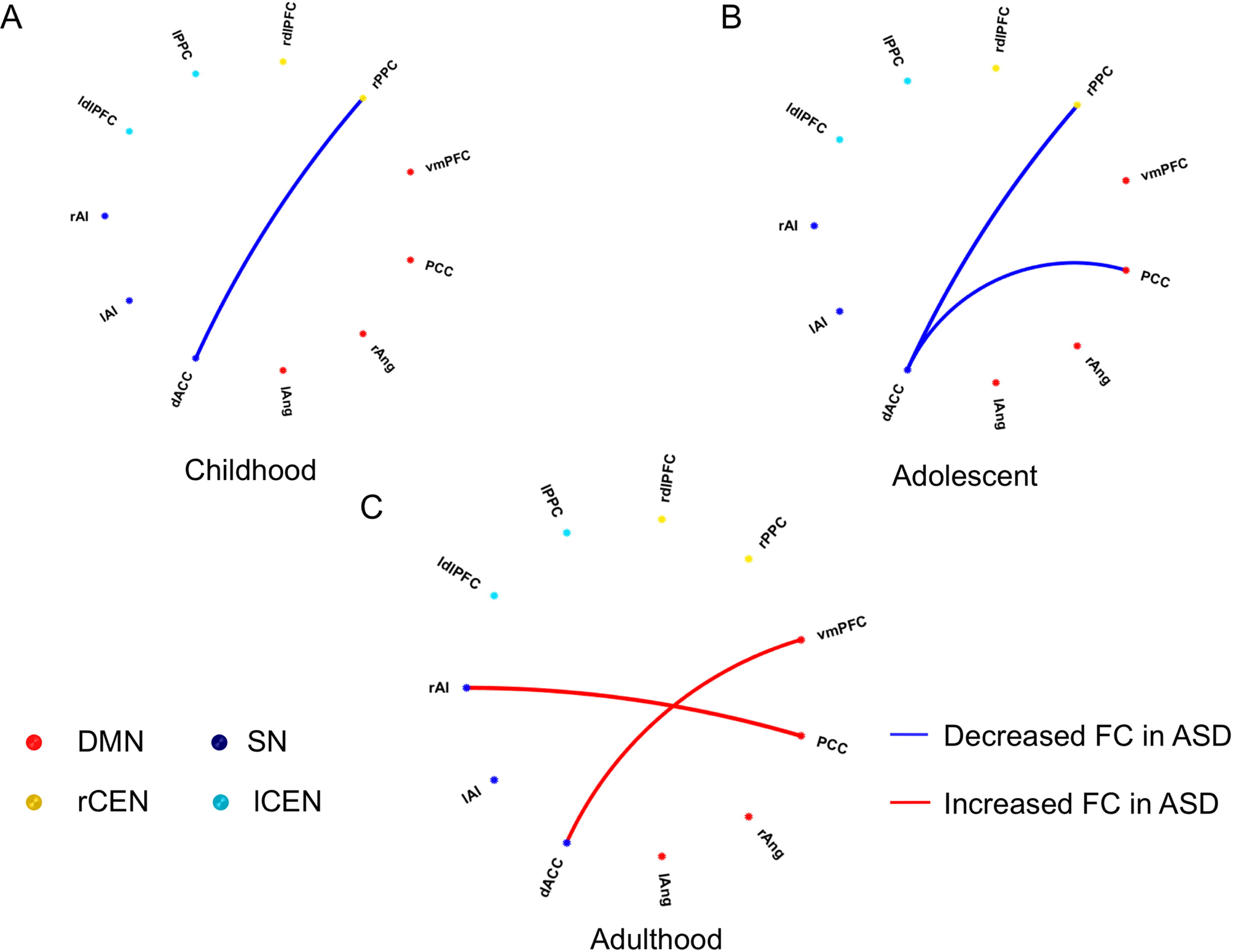
Summary of group differences in each age cohort at the level of ROI pairs’ connectivity (z). Blue lines represented decreased functional connectivity (FC) of ROI pairs in ASD, and red lines represented increased FC of ROI pairs in ASD.

### 3.3 Relationship with Symptom Severity in the ASD Group

Spearman rank-order correlation analysis was employed to explore whether the ASD symptom severity assessed by the ADOS scores was related to the indices that were identified by group differences in each age cohort. The result showed that only the connectivity of dACC-PCC and the section of communication score in adolescence were significantly related (*r*=-0.578, *p*=0.030), but this correlation did not survive after FDR correction for multiple comparisons.

### 3.4 Verification on ABIDE Dataset

To examine the robustness of our findings, we computed network-iFC using ABIDE dataset and verified the consistency in the shape of developmental trajectories of ASD and TD groups separately. In the ASD group, the shape of developmental trajectories was replicated in ABIDE. Namely, the iFC between DMN and SN remained relatively stable from age 6 to 30 years (linear: *t*=0.899, *p*=0.371; quadratic: *t*=0.959, *p*=0.340), and meanwhile the iFC between SN and rCEN increased with age (linear: *t*=2.728, *p*=0.008). However, the developmental trajectories were different between our samples and ABIDE subjects for the TD group. For DMN-SN, the iFC was unchanged across age (linear: *t*=0.073, *p*=0.942; quadratic: *t*=-0.482, *p*=0.631), and for SN-rCEN, age-related changes were revealed (linear: *t*=2.621, *p*=0.010) (**Figure 6**).

**Figure 6.**
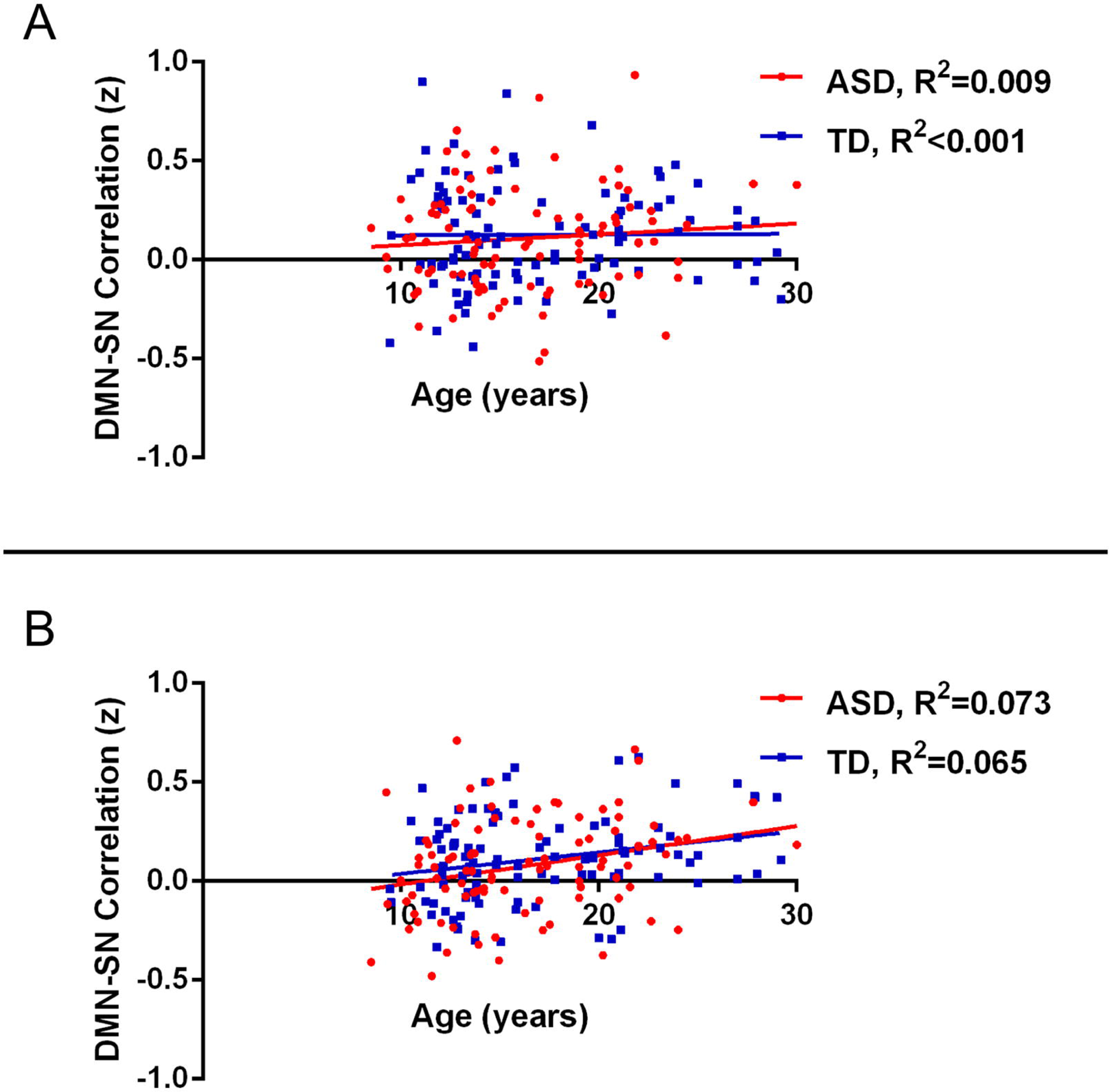
The shape of developmental trajectories at the level of network connectivity (z) using the Autism Brain Imaging Data Exchange (ABIDE) dataset. The scatter plots and fit lines showed the diagnostic group-by-age interactions (A for DMN-SN connectivity; B for SN-rCEN connectivity). The fit lines were plotted separately for the ASD (red) and TD (blue) group. DMN, SN, rCEN stand for default mode network, Salience Network and right Central Executive Network, separately.

## 4. Discussion

ASD is a heterogeneous neurodevelopmental disorder with unclear etiology and pathogenesis. The resting state fMRI has been applied to the exploration of brain mechanism in ASD for more than ten years, but results are inconsistent. The triple network model, which is related to higher cognitive function such as self, emotion, and cognitive control, has not been studied the maturation patterns of triple network iFC in ASD. To our best knowledge, this is the first study examined the functional connectivities in ASD based on the triple networks model across different age groups from childhood to early adulthood. Thus, our results enlighten a novel understanding towards the development of functional connectivity within/between the three large-scale networks (DMN, SN, rCEN, lCEN) in ASD. The ASD-specific developmental patterns and relations between aberrant brain FC and ASD symptom severity were also identified.

### 4.1 Within-network Functional Connectivity in Triple Network Model

In our study, no significant diagnostic group-by-age interactions or main effects of diagnosis in any within-network connectivity were found in the triple network model. This result indicates that the connectivity within DMN, SN, rCEN, lCEN and their maturation are intact in ASD. Our findings of the main effects of diagnosis were in line with Abbott’s research, which also revealed the result of unchanged within-network connectivity in ASD group compared to TD group (Abbott et al. 2016). However, DMN, a key network in previous studies of ASD, has been found with abnormal iFC and developmental trajectories in ASD patients; e.g., a weaker connectivity of the posterior hub with the right superior frontal gyrus, a different developmental track of the superior frontal gyrus ^39^, etc. Three factors might account for the inconsistent findings between Wiggins’s study and ours. First, only 4 ROIs were selected to represent DMN in our research, and other regions like inferior parietal lobule, lateral temporal cortex, and hippocampal formation were not included here, resulting in that only partial characteristics of within-network connectivity were explained. Second, same as Abbott et al.’s study, we took the average value of all ROI pairings within a network ^49^. The averaged value might neutralize the strength of ROI pairings’ connectivity within the networks and brought a risk of losing abnormal connectivity in ASD. Third, apart from calculation methods, the small sample size of our study is considered as an impactive factor that caused the non-significant result.

Although no significant diagnostic group-by-age interactions or main effects of diagnosis in any within-network connectivity was found we observed a linear increase of age-related iFC within DMN in both groups, which was consistent with studies of typical development ^21, 27, 35, 64–67^. The developmental characterization of “local to distributed” in functional brain networks has been proposed several years ago, which suggested that nodes with long Euclidean distance (usually over 90 mm) had weak connectivity in childhood but could increase over age ^27^. DMN is one of the networks following this principle that the average Euclidean distance between nodes (like the nodes defined in this article) is greater than 90mm. In addition, the development of DMN research confirmed sparse functional connectivity at early school age (7-9 years old), and its nodes were integrated into a cohesive network over development ^65^.

### 4.2 Between-network Functional Connectivity in Triple Network Model

Between-network results demonstrated significant diagnostic group-by-age interactions in DMN-SN and SN-rCEN. The developmental trajectories of the iFC between these networks were anomalous in the ASD group compared to the TD group. These results suggest that there are aberrant developmental patterns existing with the abnormal connectivity of the triple network model in ASD.

#### 4.2.1 The Aberrant Developmental Patterns of the Triple Network Model in the ASD Group

The predominant goal of our study was to explore age-related differences shown in triple networks for ASD patients from childhood to adulthood, which was designed as the diagnostic group-by-age interactions. We found that diagnosis-related difference in age-related changes suggests delayed, ectopic and failure, these three maturation patterns in networks’ or nodes’ iFC. As defined elsewhere ^62, 68^, delayed maturation refers to abnormalities timing of maturation (i.e., the velocity of development). On the other hand, ectopic and failure maturations imply the deviations of trajectories’ shape, which may signal more profound developmental disturbances. We also found varied developmental patterns that depend on the networks or nodes involved, which may highlight a complicated profile of age-related iFC in ASD.

Consistent with delayed maturation theory, our study revealed a relatively stable iFC between SN and rCEN from childhood to early adulthood in TD controls but not in ASD. To be specific, either at the level of networks or that of nodes (dACC-rPPC), iFCs were unchanged over the age in the TD group but were increased in the ASD group. From the view of triple-network model, the aberrant mapping from SN to rCEN gives rise to aberrant compromising cognition (e.g., working memory) and goal-relevant adaptive behaviors ^41^. A previous review summarized the impairments of executive function (e.g., response inhibition, working memory) and the compensatory mechanisms for normative function, as well as the age-related improvements from childhood to adolescence in autism ^69^. Thus, we speculate that the delayed maturation of the iFC between SN and rCEN might be the neural basis of ASD-related impairments in some aspects of executive function.

Our study also showed, either at the level of networks or that of nodes, the pattern of failure to mature was robust in the iFC between SN and DMN. In the TD group, age-related changes were both linear (SN-DMN and rAI-PCC) and quadratic (dACC-PCC), demonstrated by the iFC index, while no significant changes were found in the ASD group. Similar patterns of failure to mature have been reported in another study that examined nodes within DMN in ASD ^70^. From the perspective of triple-network model, the abnormal developmental trajectories suggested aberrant mapping between SN and DMN. Our findings displayed PCC as a locus of abnormality in ASD. The region of PCC has been typically associated with episodic memory retrieval ^71, 72^, autobiographical memory ^73^ and semantic memory related to internal thought ^74^. Part of the impairments of episodic, autobiographical and semantic memory in ASD has been established in a prior research evidenced by age-related changes of these problems ^75^. Thus, we speculate that failure in the neural maturation of iFC between SN and DMN might be the basis of ASD-related impairments in some aspects of the self-referential mental activity.

Findings of our study further showed that the developmental patterns of ASD (toward to ectopic age-related changes) accompany with delayed maturation in iFC between SN and DMN. Ectopic maturation occurs in iFC between dACC and vmPFC with an inverted U-shaped change in TD and a linear increasing change in ASD. Parallel increases in iFC strength with age in ASD have also been introduced in two previous studies; one of them focused on changes in striatal circuitry ^37^, and the other examined the functional circuitry of the posterior superior temporal sulcus from childhood to young adulthood ^38^. But problems about the influence of these abnormal brain development on clinical symptoms of ASD need to be indagated in the future.

To verify our findings, we adopted the same fMRI preprocessing and data analysis on the homogeneous samples in ABIDE I. The selected sample size from ABIDE was bigger, about double times than that collected on our own. Although the shape of development trajectories did not differ statistically between ASD and TD group, our results about the development trajectories of iFC between SN and other networks (DMN and rCEN) in the ASD group were confirmed by the analysis with ABIDE sample. Also, there were some interesting findings of the TD group in the ABIDE data that showed the opposite result of development trajectories from our data. Considering that the ABIDE data was all from the western population, further research may be necessary to explore whether an inherent discrepancy exists or not between typically developing Chinese and western populations. In addition, though the ABIDE sample size was large, there were some problems that could not be ignored, like the inconsistency in data acquisition and diagnosis caused by multi-center (e.g., the ABIDE sample was from 7 sites in this research). It might be more reliable to use public data if the “multi-center, disease heterogeneity” status could be improved in the future.

In a word, our findings demonstrated that the development of functional connectivity in triple networks in ASD is aberrant, and the aberrant development of FC in triple networks in ASD are network-specific. No single maturation pattern can be used to explain the developmental deviations of FC in triple networks in ASD. Our results propose that the factors of age, brain regions, and networks are very important for exploring brain mechanism of ASD and they should noted when interpreting group differences between ASD and TD in studies.

#### 4.2.2 The Aberrant Functional Connectivity of the Triple Network Model in the ASD Group at different ages

Based on our findings that the developmental trajectories of iFC between SN and other networks (DMN and rCEN) were aberrant in the ASD group, we divided the whole age group of ASD and TD into three cohorts of childhood, adolescence, and early adulthood to explore the group differences of iFC between SN and other networks. Accordingly, a predominant pattern of hypo-connectivity in SN-DMN or SN-rCEN was observed in children and adolescents (dACC-PCC/rPPC) but in young adults that showed hyper-connectivity (SN-DMN at the level of network, rAI-PCC, and dACC-vmPFC at the level of nodes) (**Figure 5**). These findings suggest that the ability of information exchange between SN and rCEN or DMN may be impaired in children and adolescents ASD patients, but can eventually be caught up or compensatively enhanced in early adulthood. The present findings were consistent with prior reports of within- and between-network functional connectivity in age-stratified participants with ASD and TD using the approach of independent component analyses (ICA) ^40^.

Another interesting point in our results to take note of is the atypical hemispheric asymmetries. The aberrant iFCs with rCEN and rAI suggest a shift of atypical rightward asymmetry in the ASD group. Cardinale and colleagues ^76^ have found some networks, such as visual, auditory, motor, and executive, having the same hemispheric asymmetries in ASD. Other researches also have concluded that subjects with ASD have right dominant regional homogeneity alterations of spontaneous brain activity ^76–78^. These findings corroborate the feature of atypical hemispheric asymmetry about the functional brain organization in ASD.

### 4.3 Relationship with Symptom Severity in the ASD Group

The relationship between aberrant FC with symptom severity suggests that the FC between dACC and PCC is correlated with communicative deficits in the ASD adolescents, but was absent in ASD children and adults. Weng and colleagues ^79^ also demonstrated an association of the iFC between PCC and communicative deficit in the ASD adolescents, but no association was found in ASD adults. This finding implies that the relationship between iFC and symptom severity was age-specific. However, studies with large sample size are needed to further explore the relationship between triple-network FC and symptom severity of ASD in future.

## 5. Limitations

There are some limitations need to be noted in our study. First, in terms of developmental trajectories, longitudinal study is better than cross-sectional design in dissecting inter-individual variation. A longitudinal approach would clarify the age effects or others with more sensitivity to detecting developmental changes. Second, ASD usually begins in the first 3 years of life, so the greatest developmental changes may occur earlier. Although it is difficult to operate resting state fMRI in younger children, few studies have made some attempts in toddlers with ASD ^79–81^. Third, it is likely that small sample size in each age cohorts may not be sufficient to detect correlations between iFC and ASD symptoms. Thus, large sample size, inclusion of participants under 6-year-old, and follow-up studies are needed in future work.

## 6. Conclusion

In conclusion, our study verified the aberrant of triple-network model (DMN, SN, CENs) in ASD and identified deviated development patterns of these networks. The findings are meaningful for the demonstration of network-specific and age-specific functional connectivity of ASD.

## ACKNOWLEDGEMENTS

This work was supported by the National Key R&D Program of China (2017YFC1309900, 2017YFC1309902), National Natural Science Foundation of China (81271508, 81571339, 81671774 and 81630031), Beijing Natural Science Foundation (7164314), the Hundred Talents Program of the Chinese Academy of Sciences, and Beijing Municipal Science & Technology Commission (Z161100000216152).

